# Large-scale *trans*-eQTLs affect hundreds of transcripts and mediate patterns of transcriptional co-regulation Short: *trans*-eQTLs reveal patterns of transcriptional co-regulation

**DOI:** 10.1101/056283

**Authors:** Boel Brynedal, JinMyung Choi, Towfique Raj, Robert Bjornson, Barbara E Stranger, Benjamin M Neale, Benjamin F Voight, Chris Cotsapas

## Abstract

Genetic variation affecting gene regulation is a driver of phenotypic differences between individuals and can be used to uncover how biological processes are organized in a cell. Although detecting *cis*-eQTLs is now routine, *trans*-eQTLs have proven more challenging to find due to the modest variance explained and the multiple testing burden when comparing millions of SNPs for association to thousands of transcripts. Here, we provide evidence for the existence of *trans*-eQTLs by looking for SNPs associated with the expression of multiple genes simultaneously. We find substantial evidence of *trans*-eQTLs, with an 1.8-fold enrichment in nominally significant markers in all three populations and significant overlap between results across the populations. These *trans*-eQTLs target the same genes and show the same direction of effect across populations. We define a high-confidence set of eight independent *trans*-eQTLs which are associated to multiple transcripts in all three populations, and affect the same targets in all three populations with the same direction of effect. We then show that target transcripts of *trans*-eQTLs encode proteins that interact more frequently than expected by chance, and are enriched for pathway annotations indicative of roles in basic cell homeostasis. Thus, we have demonstrated that *trans*-eQTLs can be accurately identified even in studies of limited sample size.

## Author summary

Understanding how biological processes are orchestrated requires unraveling how the genes involved are coregulated. Finding genetic variants affecting the expression of multiple genes can help identify both which genes are co-regulated and the nature of the control circuit. However, whilst mapping expression QTLs (eQTLs) close to a gene has been routine for some time, finding variants acting at a distance is more challenging as we have had to test millions of markers against thousands of transcripts. In this work we take a novel statistical approach to demonstrate the existence of *trans*-acting eQTLs that control hundreds of genes exist. The genes they control share regulatory machinery, form interaction networks and are involved in aspects of cellular homeostasis. We can thus begin unraveling the complex control architecture underlying biological processes.

## Introduction

Biological processes are carefully orchestrated events requiring precise activation and repression of participating genes by hierarchical gene regulation mechanisms. This elaborate co-regulation can be seen in the complex patterns of gene co-expression across tissues [1] and conditions [2]; the overlap and organization of transcription factor target sets [3]; the precise orchestration of developmental processes; and the organization of gene interaction networks [4]. Furthermore, it has become apparent that a substantial fraction of common genetic variants driving organismal traits such as disease risk affect gene regulatory sequences rather than coding sequence [5,6]. Thus, understanding how genetic variation influences the co-regulation of multiple genes will aid in the identification of major regulators of biological processes.

Transcript levels are heritable, with a large proportion of the variance across the human population attributable to expression quantitative trait loci (eQTLs) acting *in trans*: one recent study estimated that 88%+/−3% of transcript level heritability is due to *trans*-acting effects 7]. Unlike *cis*-acting eQTLs which are by definition localized to proximal regulatory elements [8,9], these *trans*-eQTLs are presumed to affect gene regulatory machinery encoded elsewhere in the genome [9]. Thus, a *trans*-acting variant should alter levels of all the transcripts influenced by the regulatory machinery it affects, providing a powerful way to identify co-regulated genes and eventually understand the complex, often overlapping patterns of transcriptional control [10].

Several approaches have been used to detect *trans*-eQTLs: the simplest is to treat each transcript level as an independent trait and identify regions of the genome where signals aggregate [11,12]. Genetic linkage of many transcripts to specific genomic loci in yeast [11], mouse [13,14], rat [15], maize [16] and human [16,17] suggested the presence of major *trans*-acting loci, but these have been hampered by the sensitivity of these methods to data processing artefacts [18]. In addition to thousands of *cis*-eQTLs, genetic association studies in humans have identified a limited number of *trans*-eQTLs in lymphoblasoid cell lines [19–21], adipose tissue [22,23] and whole blood samples [24]. As the majority of transcript level heritability is due to *trans*-acting influences [7], these results suggest that current eQTL cohorts lack adequate statistical power to detect *trans*-eQTLs. In particular, the correction required for the number of independent association tests across the genome *and* the number of transcripts to be analyzed imposes a heavy multiple-testing burden, whilst practical considerations limit the sample size of eQTL studies to at most several hundred individuals. An alternative approach has been to use principal component or latent variable analysis to identify trends in covariance induced by a *trans*-eQTL in the expression levels of its targets, and using this as a *meta-trait* in an association or linkage test [25,26]. This has not, to date, led to the discovery of sufficient *trans*-eQTLs to account for the 88% of heritability explained by *trans*-acting factors, indicating further approaches are warranted.

Here, we take a complementary approach to identifying *trans*-eQTLs influencing a number of transcripts. Rather than the null hypothesis of no association, the association statistic distribution at a *trans*-eQTL in the genome will be a mixture drawn from null and non-null distributions, with the proportion of non-null statistics proportional to the number of *trans*-eQTL target transcripts. We can therefore test the distribution of eQTL association statistics at each marker in the genome for evidence of deviation from the expected null (where no transcripts are associated), and infer the presence of a *trans*-eQTL if this null hypothesis is rejected (cross-phenotype meta-analysis, CPMA [27]). This second-level significance testing [28] does not identify *which* transcripts are affected by a *trans*-eQTL, but only that there exists evidence of a *trans*-eQTL. We apply this approach to publicly available eQTL data from lymphoblastic cell lines across three African HapMap populations [20], and show evidence of multiple *trans*-eQTLs in these data. We detect eight independent *trans*-eQTLs associated with multiple transcripts in all three populations, and where the transcript targets overlap significantly and the direction of effect is the same across the three populations. We then show that target transcripts of *trans*-eQTLs encode proteins that interact more frequently than expected by chance, and are enriched for pathway annotations indicative of roles in basic cell homeostasis, suggesting they are co-regulated sets of genes.

## Methods

Unless otherwise stated, all statistical analyses were done using the R programming language (v 3.1.0) [29]. Additional libraries are cited where appropriate. An overview of our pipeline is shown in Figure S1 and our pipeline is available for download at [[http://www.github.com/cotsapaslab/]].

### Genotype data processing

We selected to study unrelated individuals from the three African populations included in the HapMap project phase III, reasoning that the high genetic diversity and average minor allele frequencies observed in Africa will increase the statistical power of the eQTL association tests. We obtained genome-wide genotype data for 135 Maasai in Kinyawa, Kenya (MKK); 83 Luhya in Webuye, Kenya (LWK); and 107 Yoruba in Ibadan Nigeria (YRI) from the HapMap Project website (ftp://ftp.ncbi.nlm.nih.gov/hapmap/genotypes/2009-01_phaseIII/plink_format/; accessed: 2014-06-18). As our sample size is limited, we restricted our analysis to 737,867 autosomal markers with at least 15% minor allele frequency in all three populations. All remaining variants are in Hardy-Weinberg equilibrium (*P_HWE_ > 1e^-6^*); all individuals have <3% of genotypes missing and all remaining variants have <8% missing data. Genotype data annotation was converted into hg38 coordinates.

### Expression data processing

We obtained processed expression data for lymphoblastoid cell line profiling on the Illumina Human-6 v2 Expression BeadChip array for all 322 individuals, publicly available under ArrayExpress accession number E-MTAB-264 [20]. The expression data includes 21,802 probes mapping to one single gene, excluding probes that map to multiple genes or to genes on the X or Y chromosome, and that have not been subjected to the PEER method [20]. After quantile normalization to reduce inter-individual variability [30], we removed probesets with low variance or low intensity in each population. Both the interquartile range and mean intensity across probe sets showed clear bimodal shapes (Figure S1), and we used mixture modeling (mclust v.5.1 in R) to detect those probe sets that belonged to each higher distribution with 80% probability. We retained those probe sets that had a higher variance and higher intensity in all three populations, resulting in 9085 analyzed probe sets.

By converting Illumina probe IDs to HGNC gene symbols (biomaRt v2.22.0 [30]) we could map 8673/9085 probesets to 7984 unique HGNC genes with unambiguous hg38 genomic coordinates in GENCODE v.20. Unmapped probesets were excluded from analyses relying on annotation.

Expression data suffer from systematic, non-genetic biases, hampering eQTL studies [18]. Several multivariate approaches have been used to correct these data artefacts [25,31,32], all of which identify trends in variance in expression data assumed to stem from (usually unmeasured) confounders. These methods clearly improve power to detect *cis*-eTQLs [23,33], but cannot distinguish between systematic artefacts and genuine *trans*-eQTLs, both of which will explain some proportion of variance across many transcripts [25,31,32]. For this reason, we have chosen not to use these corrections in our data processing pipeline, as our goal is to detect the presence of *trans*-eQTLs.

### Calculating eQTL association statistics

We calculated association statistics for each probeset intensity to each SNP by linear regression [34], controlling for population stratification by adding structure principal components as covariates [35]. In each population, we estimated the optimal number of principal components by incremental inclusion of components until the overall test statistic inflation is minimized, as previously described [36] (see Supplementary Information). We included the top two principal components for YRI, ten for LWK and 20 for MKK as optimal corrections for population stratification.

### Identifying *trans*-eQTLs by cross phenotype meta-analysis

Previous strategies to identify *trans*-eQTLs rely on either identifying significant associations to a single transcript [37,38], or associating variance components affecting multiple transcripts with genetic markers as surrogate phenotypes [25,39]. We have previously described a second-level significance testing approach [28] to assess evidence of multiple associations at a genomic marker [27]. At each marker we test for overdispersion of association −*log(p)* values across all probe sets with a null hypothesis that −*log(p)* should be exponentially distributed, with a decay parameter *λ* = 1. Under the joint alternative hypothesis, where a subset of association statistics are non-null, *λ ≠* 1. We compare the evidence for these hypotheses as a likelihood ratio test for our cross-phenotype meta-analysis (CPMA), where the statistic *S_CPMA_* is defined as:

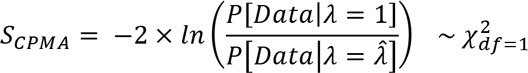

where 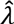 is the observed exponential decay rate in the data. Thus we need only estimate a single parameter, 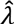, so that the test has a single degree of freedom.

We account for the extensive correlation between *t* probeset levels across individuals by empirical significance testing. We simulate eQTL association statistics under the null expectation of no association to any marker given the observed correlation between probe sets association statistics from a multivariate normal distribution (using the MASS package in R [40]). We perform an eigen-decomposition:

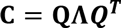

Where the covariance matrix **C** has entries *c_i,j_* = *cov*(*a_i_,a_j_*) where *a_i_* and *a_j_* are vectors of scaled z-scores for the *i*th and *j*th probesets across all markers in the genome. All three sample covariance matrices thus have dimension 9085; because they are calculated from the probeset x SNP matrix of eQTL Z statistics rather than the probeset x individual matrix of expression levels, we find all three are positive definite (data not shown). To account for the correlation between transcript expression levels, we generate the empirical null distribution *Z*\* of association statistics using:

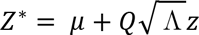

Where *z* is a vector of i.i.d. standard normal values *(N(0,1))* and *μ* a vector of mean eQTL Z statistics of the 9085 probe sets.

We calculate *p* values from this null distribution, calculate *S_CPMA_* and determine empirical significance as:

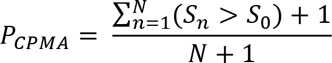

where *S_n_* is *S_CPMA_* for the *n*th iteration of the null simulation, *S_0_* is the observed *S_CPMA_* and *N* is the number of permutations (here, N = *5,000,000*).

We investigate the overlap of signal across populations using hypergeometric tests (hint v 0.1-1) of the independent SNPs (see below) at FDR α levels 0.5, 0.4, 0.3, 0.2, 0.1 and 0.05 [41]. We also investigate the enrichment across populations using Wilcoxon sum-rank test at different α levels.

### Meta-analysis of CPMA statistics

We combined empirical CPMA statistics from the three African populations using sample-size weighted metaanalysis [42]. To identify independent effects across the genome, we clumped these meta-analysis results at r^2^ < 0.2 [34].

### Analytical validation of *trans*-eQTLs

To validate the detected *trans*-eQTLs, we perform two secondary analyses: we test if the *trans*-eQTL is associated to the same probesets in the three populations, and if the directions of effect are consistent across the three populations. This information is not used in the CPMA and meta-analysis calculations, and thus offers an independent validation analysis on these data.

We first empirically assess evidence that a *trans*-eQTL is associated to the same probesets across populations. In a pair of populations *P_1_* and *P_2_*, we observe *N_1_* and *N_2_*) probesets with an eQTL *p <= 0.05* at a *trans*-eQTL, with an intersect *N_0_*=*(N_1_* ∩ *N_2_)*. We construct the expected distribution of *N_o_* using the *N_1_* and *N_2_* most associated probesets at all *M* independent SNPs across the autosomes, and compute empirical significance *P_0_* as:

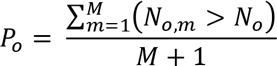

Similarly, we assess consistency of effect direction for *N_O_* probesets with an association *p* <= *0.05* to a *trans*-eQTL in a pair of populations P_1_ and P_2_. To allow for alternate linkage disequilibrium patterns in different populations (where effects can be opposite with respect to a detected *trans*-eQTL) we define the overlap in directionality as *N_dir_* = *max((N_1,p_* ∩ *N_2,p_)* ∪ *(N_1,n_* ∩ *N_2,n_)*, *(N_1,p_* ∩ *N_2,n_)* ∪ *(N_1,n_* ∩ *N_2,p_))* where *N_1,p_* are the number of probe sets with increasing expression given the number of alleles of the SNP, and *N_1_,_n_* those with decreasing expression. We construct the null distribution of *N_dir_* of the targets of each *trans*-eQTL by computing it for all *M* independent SNPs across the autosomes and compute empirical significance *P_dir_* as:

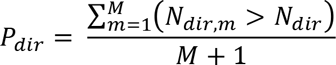

We used hypergeometric tests to assess the significance of the intersections between all three populations [41] (R package hint v. 0.1-1).

### Defining high-confidence *trans*-eQTL targets

We used a meta-analytic approach to define consensus target gene sets for the ten high-confidence *trans*-eQTLs. For each candidate *trans*-eQTL, we meta-analyzed eQTL association statistics for each of the 9,085 probesets across the three populations using sample-size weighted fixed effect meta-analysis [34,42], and then defined the group of target probesets as those with FDR < 0.01. This approach differs from the metaanalysis of the aggregate CPMA statistics above, where we are combining overall evidence of a *trans*-eQTL rather than for association to specific probeset levels.

### Functional enrichment analyses of *trans*-eQTL target probesets

For each set of *trans*-eQTL target transcripts, we calculated enrichment of proximal transcription factor binding events using publicly available chromatin immunoprecipitation/sequencing (ChIPseq) data for 50 factors in lymphoblastoid cell lines from the ENCODE consortium [3,43]. We were able to annotate 2405/7984 unique HGNC genes corresponding to the 9085 probesets in our analysis with at least one transcription factor binding event from these data. We observed *TF_o_*, the number of binding events for each transcription factor in the target probesets of each *trans*-eQTL, and assessed significance empirically by resampling probesets with similar expression intensity over *N=1,000* iterations:

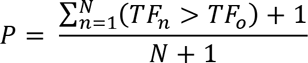

where *TF_n_* is the number of binding events for a transcription factor in the *n*th iteration.

To test for functional categories over-represented in each set of *trans*-eQTL target transcripts, we looked for enrichment of Gene Ontology biological process annotations with the hypergeometric approach implemented in BioConductor [30], which accounts for the dependencies in the hierarchical structure of the ontology. We only considered terms where at least 10 genes were observed.

To establish if each set of *trans*-eQTL target transcripts represent biological networks we used our previously described **P**rotein **I**nteraction **N**etwork **T**issue **S**earch (PINTS) framework [44] (R packages PINTS v. 0.1, igraph v. 1.44.1 and BioNet v. 1.29.1). Briefly, for each *trans*-eQTL we first collapse target probesets onto HGNC genes, and then project these onto a protein-protein interaction network. We detect the largest subnetwork of target genes using the prize-collecting Steiner tree algorithm, and assess significance by permuting the network 100 times and assessing the size and connectivity of the largest subnetwork in the observed data. For any subnetworks showing significant excess in either size or connectivity, PINTS then tests for preferential expression across a tissue atlas [44].

## Results

### Replicable *trans*-eQTLs affect many genes

We sought evidence of *trans*-eQTLs affecting the expression levels of many target genes by assessing if there is excess eQTL association at common autosomal variants compared to chance expectation [27] across three African HapMap populations [20]. We analyzed population structure-corrected eQTL data for 9085 probe sets at 737,867 autosomal markers from the MKK, LWK and YRI HapMap populations (135, 83 and 107 individuals respectively [20]), empirically assessing significance to account for the correlation between eQTL statistics. We first compared the three cohort analyses to assess consistency in marker statistics indicative of replication, and find overlap between all three populations (Table 1 and S1) suggesting the presence of true *trans*-eQTL. To further explore these results, we meta-analyzed our CPMA statistics across the three cohorts (Supplementary Figure 5), and found 16,484/178,464 (9.2%) pairwise-independent SNPs with meta-analysis *p_meta_* < 0.05, though none reached genome-wide significance (minimum *p_meta_* = 7.2 × 10^−7^ at rs10842750).

**Table 1.** CPMA statistics are consistent across populations, indicating the presence of true *trans*-eQTLs. We observe strong overlap between variants with modest CPMA statistics across the three populations, indicating the presence of many *trans*-eQTL effects across the genome. Our results indicate limited power to detect any single *trans*-eQTL, likely due to limited sample size, suggesting that substantially larger sample sizes than the 322 in our current dataset will be required to power discovery.

| CPMA $\alpha$ | Expected | Observed | *Hypergeometric*<br>*P* value |
| --- | --- | --- | --- |
| 0.5 | 37,942 | 39,479 | $3.6 \times 10^{-40}$ |
| 0.4 | 20,459 | 21,361 | $1.7 \times 10^{-18}$ |
| 0.3 | 9,048 | 9,350 | $8.2 \times 10^{-5}$ |
| 0.2 | 2,815 | 2,787 | 0.73 |
| 0.1 | 362 | 361 | 0.54 |

We next sought to prioritize high-confidence *trans*-eQTLs from the 16,484 candidates with additional independent criteria, which CPMA does not consider. We expect true *trans*-eQTLs to fulfill two predictions: the genes they influence should be the same across populations; and the direction of effect should be consistent between the populations for these genes. A major technical issue to testing these predictions is the extensive correlation between gene expression levels (and therefore between eQTL association statistics), so we assess significance for both these predictions empirically (see methods) in pairwise comparisons of populations. Of the 16,484 *trans*-eQTLs, we found that 1,692 (10.2%; YRI and MKK), 1,851 (11.2%; YRI and LWK) and 1,892 (11.5%; MKK and LWK) show nominal significance of target overlap (empirical overlap *P < 0.05;* Figure 1). Furthermore, 62 *trans*-eQTLs have significant target overlaps across all three pairwise comparisons (22 overlaps expected by chance, hypergeometric *p* = 4.5 × 10^−13^). This suggests the presence of multiple *trans*-eQTLs affecting the same target genes across populations.

**Figure 1.**
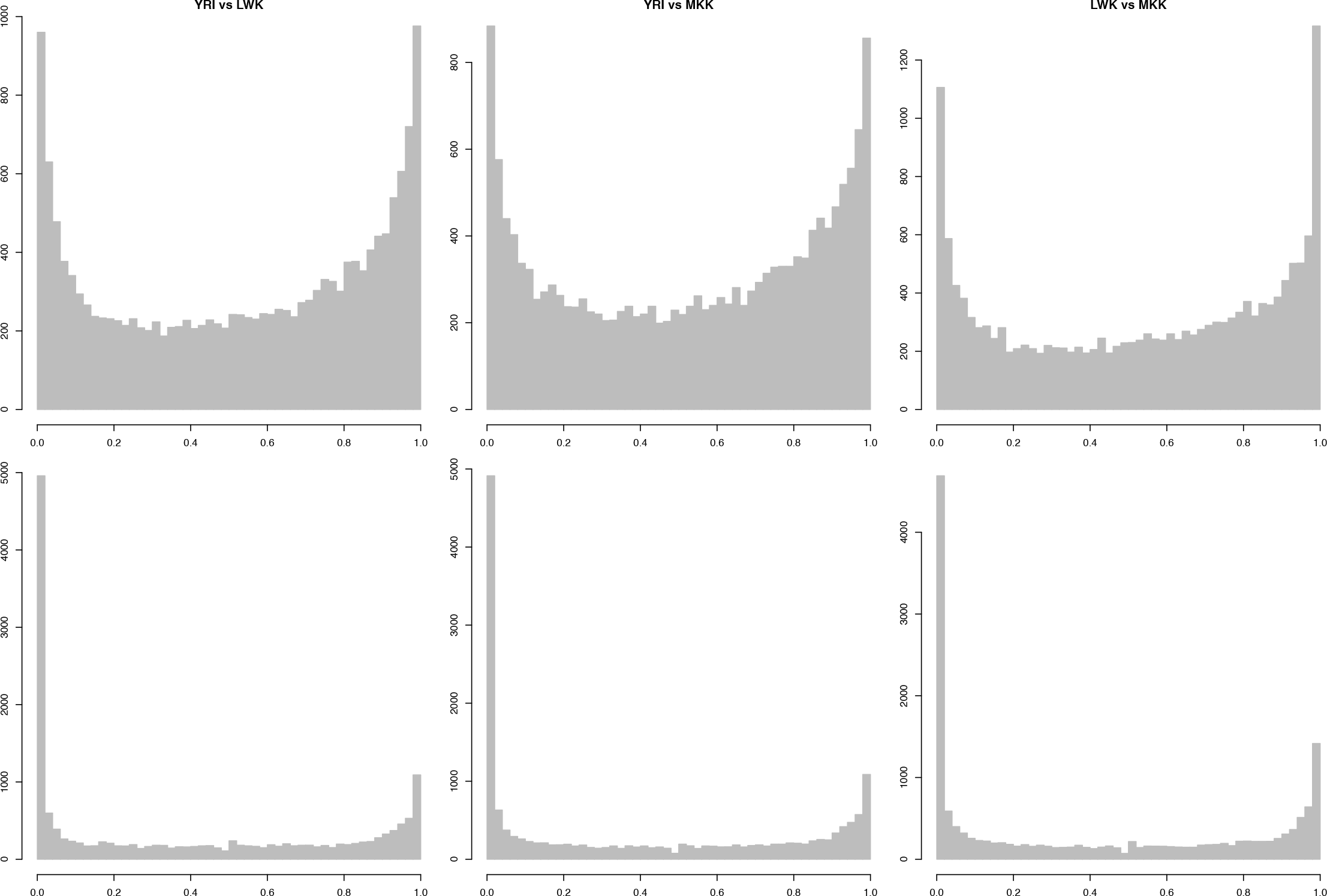
Hundreds of putative *trans*-eQTLs across the genome affect the same genes in the same direction across three African Hapmap populations. We considered all autosomal variants with nominal evidence of association to multiple transcript levels (*P_cpma_* < *0.05*). We find that they tend to target the same transcripts (top, empirical assessment of *trans*-eQTL target overlap between pairs of populations); and that the allelic effects are consistently in the same direction (bottom, empirical assessment of *trans*-eQTL sign tests between pairs of populations).

To test our second prediction, we sought evidence that the direction of effect is consistent across two populations. We find that 5,743 (34.8%; YRI and MKK), 5,762 (35.0%; YRI and LWK) and 5,498 (33.4%; MKK and LWK) of 16,484 candidate *trans*-eQTLs show nominal significance for consistent effects, and it is the same *trans*-eQTLs generating these signals (Figure 1). Furthermore, 1,062 *trans*-eQTLs are significant across all three pairwise comparisons (670 overlaps expected by chance, hypergeometric *p* = *8.1* × *10^−64^*). We also find the target overlap and directionality overlap statistics are significantly correlated (Figure S6 *p* < 2.2 × 10^−16^), indicating the presence of *trans*-eQTLs affecting the same target transcripts in the same way. Thus, our results provide several lines of evidence for *trans*-eQTLs replicating across multiple populations.

### Target genes of *trans*-eQTLs are co-regulated

We identified a high-confidence set of ten *trans*-eQTLs that are nominally significant in our CPMA metaanalysis and in all the above pairwise tests of target overlap and directionality, including two with a small number of targets, which we excluded from further consideration (Table 2). For each *trans*-eQTL, we defined a consensus set of target transcripts (FDR < 0.01) by meta-analyzing eQTL statistics for individual probesets across the three populations, rather than defining consensus sets by overlapping putative target lists at arbitrary thresholds from the individual populations (Figures 2, 3 and Supplementary Figures 7A-F).

**Figure 2.**
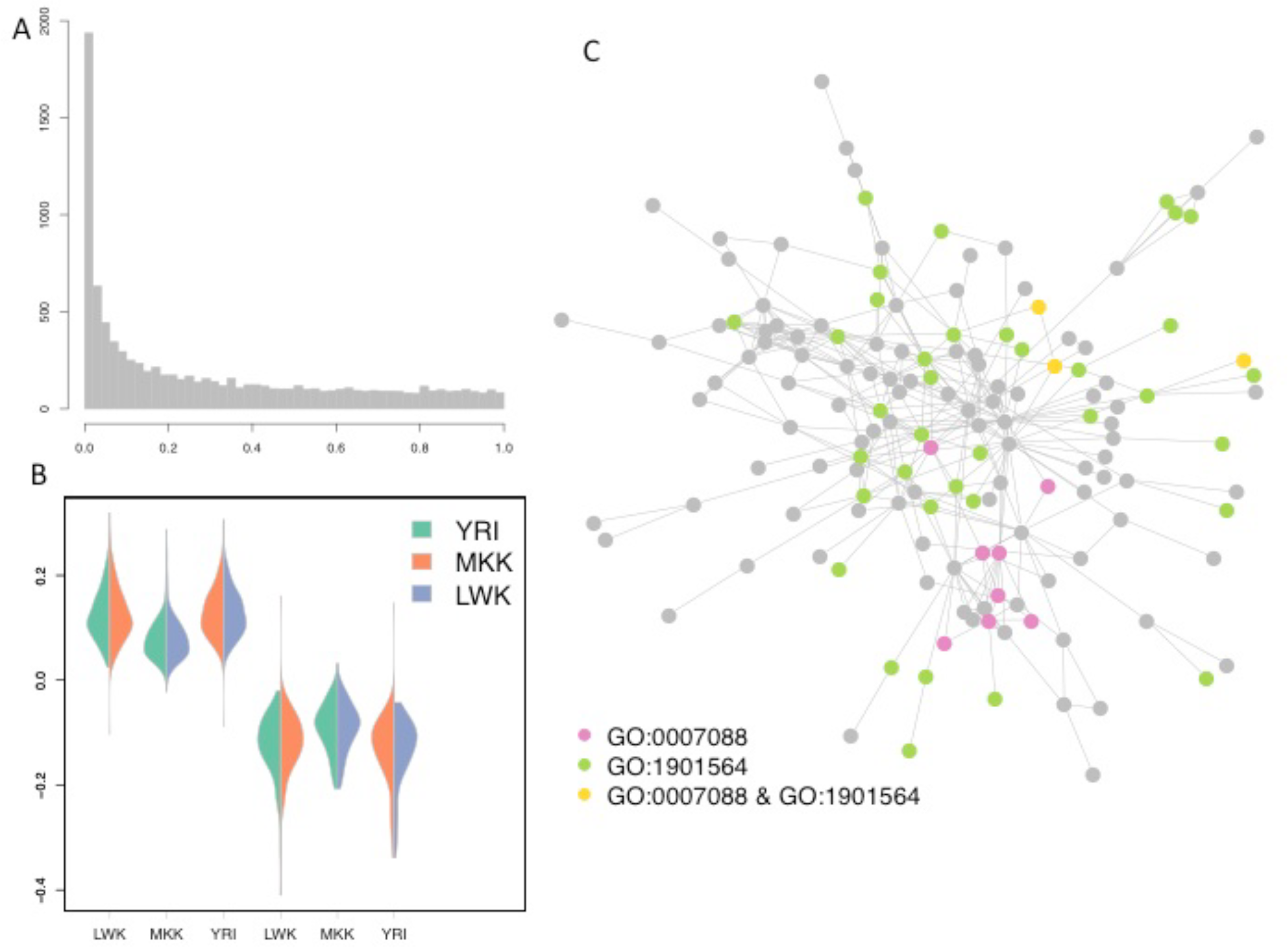
A *trans*-eQTL at rs6899963 on chromosome 6 affects the expression levels of many genes across three African HapMap populations. (A) Meta-analysis *p*-values for 9,085 transcript eQTLs at rs6899963. (B) Effect directions are consistent across the three populations. In each population (x axis), we select SNPs where the minor allele increases (left) and decreases (right) expression, respectively, and show the direction of effect in the other two populations as violin plots. The overwhelming majority of effects are consistent across all three populations. (C) The target genes of the rs6899963 *trans*-eQTL form a large subnetwork, which is enriched for multiple Gene Ontology biological processes. Here, we show the interplay between the top two enriched terms: GO:0007088 (*regulation of mitotic nuclear division*) and GO:1901564 (*organonitrogen compound metabolic process*).

We predict that if the target transcripts of our eight high-confidence *trans*-eQTL are co-regulated, they should represent a limited number of biological pathways. We therefore looked for enrichment of transcription factor binding events upstream of each of the eight target gene groups, using chromatin immunoprecipitation and sequencing (ChlPseq) data from lymphoblasoid cell lines profiled by the ENCODE project [3,43]. We found significant enrichment for at least one transcription factor in four out of the eight *trans*-eQTLs (Table 2), suggesting that *trans*-eQTL target genes are regulated by the same cellular mechanisms.

**Table 2.** Eight *trans*-eQTLs affect hundreds of transcripts across the genome. We identified a subset of *trans*-eQTLs with nominally significant CPMA meta-analysis statistics^a^; pairwise tests of target overlap^b^; and pairwise tests of directionality^c^ across three populations. For each *trans*-eQTL, we defined a consensus set of target transcripts^d^ (FDR < 0.01) by meta-analyzing eQTL statistics for individual probesets across the three populations, and find significant enrichment of transcription factor binding events at their promoters^e^. These targets also form significant protein-protein interaction subnetworks^f^.

| *trans*-eQTL | position | Within gene | CPMA<sup>a</sup><br>($P_{meta}$) | Target overlap<sup>b</sup><br>($P$) | Effect direction<sup>c</sup><br>($P$) | Target genes<sup>d</sup> | TF motifs enriched<sup>e</sup> | Network enrichment<sup>f</sup><br>($P$) |
| --- | --- | --- | --- | --- | --- | --- | --- | --- |
| **rs7694213** | chr4: 53157184 | SCFD2 | $1.6 \times 10^{-2}$ | $1.7 \times 10^{-4}$<br>$4.0 \times 10^{-2}$<br>$3.1 \times 10^{-3}$ | $5.6 \times 10^{-6}$<br>$2.0 \times 10^{-4}$<br>$5.6 \times 10^{-6}$ | 417 | | 0.18 |
| **rs6899963** | chr6: 104048696 | | $1.9 \times 10^{-2}$ | $3.5 \times 10^{-2}$<br>$5.1 \times 10^{-3}$<br>$2.0 \times 10^{-2}$ | $5.6 \times 10^{-6}$<br>$1.7 \times 10^{-5}$<br>$3.9 \times 10^{-5}$ | 393 | SRF, STAT3 | $<10^{-3}$ |
| **rs9406332** | chr6: 169317960 | | $3.7 \times 10^{-2}$ | $3.4 \times 10^{-2}$<br>$2.8 \times 10^{-2}$<br>$3.9 \times 10^{-2}$ | $3.4 \times 10^{-5}$<br>$1.1 \times 10^{-5}$<br>$5.3 \times 10^{-4}$ | 77 | | 0.21 |
| **rs10107976** | chr8: 62689355 | NKAIN3 | $7.9 \times 10^{-3}$ | $1.7 \times 10^{-3}$<br>$3.4 \times 10^{-2}$<br>$7.7 \times 10^{-3}$ | $1.7 \times 10^{-5}$<br>$5.6 \times 10^{-6}$<br>$5.6 \times 10^{-6}$ | 116 | USF1, USF2,<br>MAX, RFX5,<br>ZZZ3 | 0.19 |
| **rs4773419** | chr13: 111658732 | RP11-65D24.2 | $2.4 \times 10^{-2}$ | $5.6 \times 10^{-3}$<br>$2.6 \times 10^{-2}$<br>$1.3 \times 10^{-2}$ | $5.6 \times 10^{-6}$<br>$1.2 \times 10^{-4}$<br>$1.2 \times 10^{-4}$ | 166 | SP1, PBX3,<br>FOS, NRF1,<br>BRCA1 | 0.37 |
| **rs11621120** | chr14: 29324177 | RP11-562L8.1 | $1.9 \times 10^{-2}$ | $8.5 \times 10^{-4}$<br>$8.2 \times 10^{-3}$<br>$4.9 \times 10^{-3}$ | $5.6 \times 10^{-6}$<br>$5.7 \times 10^{-4}$<br>$7.8 \times 10^{-5}$ | 228 | | 0.19 |
| **rs10520643** | chr15: 86859175 | AGBL1 | $2.4 \times 10^{-3}$ | $4.5 \times 10^{-3}$<br>$2.3 \times 10^{-2}$<br>$1.3 \times 10^{-2}$ | $5.6 \times 10^{-6}$<br>$5.6 \times 10^{-6}$<br>$3.9 \times 10^{-5}$ | 833 | NR2C2 | $<10^{-3}$ |
| **rs7281608** | chr21: 23675283 | | $4.5 \times 10^{-2}$ | $2.8 \times 10^{-2}$<br>$4.8 \times 10^{-3}$<br>$7.1 \times 10^{-3}$ | $1.5 \times 10^{-2}$<br>$1.1 \times 10^{-2}$<br>$4.5 \times 10^{-4}$ | 97 | | $1 \times 10^{-2}$ |

We next assessed pathway enrichment using of Gene Ontology annotations and found strong enrichment for all eight *trans*-eQTL gene target sets, with 20-100 biological processes significant in each set (Table S2 lists the 10 most significant biological processes of each *trans*-eQTL). Notably, we find that target sets are enriched for fundamental biological processes including cell cycle control, metabolism and assembly of cellular machinery. To further characterize these functional connections, we assessed if the target gene sets form interacting protein networks, and find that three out of the eight *trans*-eQTL target sets form interaction networks larger and more densely connected than expected by chance [44] (Table 2). We also find that these subnetworks are preferentially expressed in particular tissues: the largest subnetwork of rs6899963 target genes (network permutation tests: size *P* = 0.03; number of edges *P* < 0.001; connectivity coefficient *P* < 0.001; overall eQTL statistic load *P* = 0.02; Figure 2) is preferentially expressed in fetal tissues and inducible pluripotent stem cells. The largest subnetwork of rs10520643 target genes (network permutation tests: size *P* = 0. 23; number of edges *P* < 0.001; connectivity coefficient *P* < 0.001; overall eQTL statistic load *P* = 0.18; Figure 3), is preferentially expressed in a similar pattern across fetal tissues and inducible pluripotent stem cells. Collectively, these results show that *trans*-acting eQTLs modulate transcriptionally coherent groups of genes involved in basic cellular processes.

**Figure 3.**
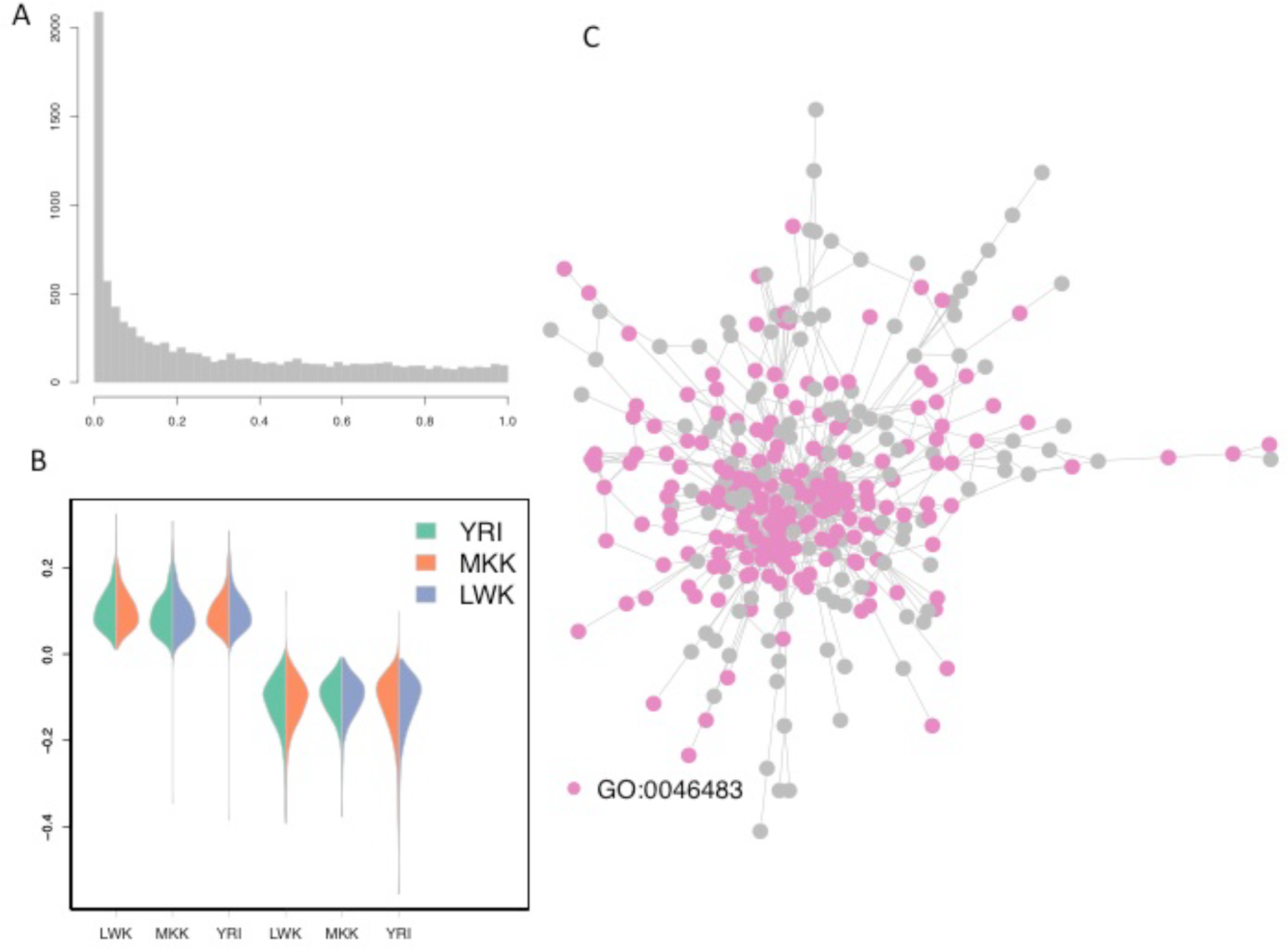
A *trans*-eQTL at rs10520643 on chromosome 15 affects the expression levels of many genes across three African HapMap populations. A. Meta-analysis p-values for 9,085 transcript eQTLs at rs10520643. B. Effect directions are consistent across the three populations. In each population (x axis), we select SNPs where the minor allele increases (left) and decreases (right) expression, respectively, and show the direction of effect in the other two populations as violin plots. The overwhelming majority of effects are consistent across all three populations. C. The target genes of the rs10520643 *trans*-eQTL form a large subnetwork, which is enriched for multiple Gene Ontology biological processes. Here, we show the term GO:0046483 (*heterocycle metabolic process*).

## Discussion

In this work we present evidence for *trans*-eQTLs by identifying SNPs simultaneously associated to the levels of many transcripts. We are able to show that these replicate across populations, and associate to the same genes in the same direction. Furthermore, we show that the target gene sets for eight high-confidence *trans*-eQTLs are bound by the same transcription factors, are enriched for pathway annotations and form significant interacting networks. Thus we conclude that *trans*-eQTLs can be identified even in studies of limited sample size.

*Trans*-eQTLs have proven challenging to detect in human data, despite the substantial heritability of gene expression attributed to them [7]. This difficulty is driven both by the modest effect sizes of *trans*-acting variants [21,23] and the systematic noise in gene expression assays [18]. Whilst both issues can be addressed by increasing sample size to boost statistical power [45] and the emergence of more technically robust assays like RNA sequencing [37], the cost and logistics of ascertaining large cohorts remains economically daunting, especially when considering multiple tissues [46]. Our approach, like many other novel analytical methods [25,32], can help maximize the insights gleaned from current resources.

We note that our approach is geared towards detecting *trans*-eQTLs influencing many genes, at the cost of low power to detect effects on single genes or a small number of targets [22,23,45]. However, larger sample sizes will be required to estimate the relative contributions of *trans*-eQTLs affecting many genes and those affecting few targets to the overall heritability of transcript levels, so we cannot yet gauge how widespread variation in large transcriptional control networks may be. Our results are, however, consistent with precise regulation of biological processes at such a large scale, particularly for basic homeostatic mechanisms. These observations further support the notion that regulation of basic cell processes is highly orchestrated and occurs on several levels simultaneously [47]. Applying this approach to eQTL datasets from diverse tissues under different stimuli will yield rich insights into tissue-specific regulatory circuits driving diverse cellular processes. Finally, we note that biological exploration and dissection of these pathways will require new experimental tools, which can address the subtleties of quantitative regulatory changes in large numbers of genes.

## Acknowledgements

Computing resources at Yale were funded partly by NIH grants RR19895 and RR029676-01. BB was supported by a post-doctoral fellowship from the Swedish Research Council.

